# Non-motile subpopulations of *Pseudomonas aeruginosa* repress flagellar motility in motile cells through a type IV pili- and Pel-dependent mechanism

**DOI:** 10.1101/2021.10.21.465385

**Authors:** Kimberley A. Lewis, Danielle M. Vermilyea, Shanice S. Webster, Christopher J. Geiger, Jaime de Anda, Gerard C. L. Wong, George A. O’Toole, Deborah A. Hogan

## Abstract

The downregulation of *Pseudomonas aeruginosa* flagellar motility is a key event in biofilm formation, host-colonization, and the formation of microbial communities, but the external factors that repress motility are not well understood. Here, we report that under swarming conditions, swarming motility can be repressed by cells that are non-motile due to the absence of a flagellum or flagellar rotation. Non-swarming cells, due to mutations that prevent either flagellum biosynthesis or rotation, present at 5% of the total population suppressed swarming of wild-type cells under the conditions tested in this study. Non-swarming cells required functional type IV pili and the ability to produce Pel exopolysaccharide to suppress swarming by the flagellated wild type. In contrast, flagellated cells required only type IV pili, but not Pel production, in order for swarming to be repressed by non-flagellated cells. We hypothesize that interactions between motile and non-motile cells may enhance the formation of sessile communities including those involving multiple genotypes, phenotypically-diverse cells, and perhaps other species.

**Importance:** Our study shows that, under the conditions tested, a small population of non-swarming cells can impact the motility behavior of the larger population. The interactions that lead to the suppression of swarming motility require type IV pili and a secreted polysaccharide, two factors with known roles in biofilm formation. These data suggest that interactions between motile and non-motile cells may enhance the transition to sessile growth in populations and promote interactions between cells with different genotypes.

## Introduction

Microbial localization through processes other than cell-division are critical for the formation of spatially structured populations and communities. Thus, motility and its regulation by diverse chemical and physical stimuli are major drivers of intraspecies and interspecies microbial interactions (1, 2). Previously, we found that exogenous ethanol, a common product secreted by many microbes, markedly reduced *Pseudomonas aeruginosa* swimming and swarming motilities at the population level, but only decreased the motile fraction of cells in the population by 16% (3). Prompted by this finding, we sought to determine the impact of a subpopulation of non-motile cells on the motility of the larger motile *P. aeruginosa* population.

During *P. aeruginosa* biofilm formation, motile cells are recruited to microcolonies of sessile cells via a combination of processes that may differ between strains with distinct early-biofilm forming strategies (4, 5) or in response to different experimental conditions. For *P. aeruginosa* strain PA14, retractile type IV pili (T4P) participate in surface sensing, which results in the upregulation of cAMP and the subsequent induction of cyclic-di-GMP (6-8); flagellar motility is then down-regulated in order to facilitate surface attachment (6, 7, 9, 10) followed by Pel exopolysaccharide matrix production (11), which can connect cells to one another (10, 12). Both matrix production and T4P function have been shown to participate in motility repression (10, 12-14) suggesting that matrix materials may mediate physical interactions between neighboring cells. T4P also mediate cell-cell interactions in swarms (15).

Boyle et al. (16) previously showed that a Δ*flgK* mutant lacking a flagellum repressed swarming (i.e., decreased the swarm radius) when in excess (83%) of the flagellated wild type (WT) (17%). In support of the observation that a non-motile subpopulation can suppress population-wide swarming, our previously published findings (3) showed that while WT strain PA14 was capable of swarming, an average of only 38% of flagellated cells were motile at any given time. Furthermore, we showed that ethanol strongly repressed *P. aeruginosa* swarming motility by reducing the motile fraction of the population from 38% to 22%; in other words, a 16% reduction in the motile fraction was sufficient to repress swarming by the entire population (3). Taken together, these findings show that a non-motile subpopulation is capable of repressing swarming in the entire population. However, how non-motile cells, such as those lacking a flagellum, suppress swarming of flagellated strains has not been elucidated.

In this study, we show that non-motile cells lacking a flagellum, when added to a population at 5 to 75% of the total inoculum, resulted in repression of swarming by WT cells. To repress flagellar motility, non-motile cells required the ability to produce Pel polysaccharide, and both motile and non-motile cells required functional T4P for this interaction. These data have implications for factors that contribute to population-level behaviors and intra- and inter-species interactions. Interactions that promote cell-cell interactions may be relevant in situations where non-motile and motile cells can be found together such as in cystic fibrosis (CF)-associated lung infections (17-19), wounds (20), and ear infections (18).

## Results

### The presence of non-flagellated cells represses swarming motility by *P. aeruginosa* strain PA14

To explore the effects of a subpopulation of non-motile cells on swarming motility, mixes containing different proportions of flagellated *P. aeruginosa* strain PA14 (WT) cells and non-flagellated Δ*flgK* mutant cells that lack the flagellar hook protein were plated on 0.5% M8 agar, which supports swarming motility by the WT strain. The percent of Δ*flgK* mutant cells ranged from 5% to 75% in these mixes, and single strains were included as controls. As expected, cultures with 100% WT swarmed readily while cultures with 100% Δ*flgK* showed no swarming (**Fig. 1A**). We found that swarming motility was completely repressed by the addition of 5-75% Δ*flgK* cells under the conditions tested (**Fig. 1A** and **Fig. 1C**). The addition of 1-4% Δ*flgK* was insufficient to repress population-wide swarming (**Fig. S1**). It is worth noting that a swarming phenotype can be variable between both technical replicates and biological replicates (21, 22). While Δ*flgK* added at 5% of the population regularly repressed WT swarming, the lower limit was variable (**Fig. S1, Fig. S5**). However, work by Boyle et al. (16) reported that Δ*flgK* repressed swarming only when it was in excess of the WT. Specifically, the authors reported repression as measured by a reduction in swarm radius when there was 17% WT and 83% Δ*flgK*. Images for 1:1 WT:Δ*flgK* mixes may also suggest effects on the swarm area even though swarm radius was not yet affected. Therefore, while the lower limit of our assay was variable, Δ*flgK* repressed WT swarming at significantly lower concentrations than previously reported and did not depend on being in excess of the WT.

**Fig. 1.**
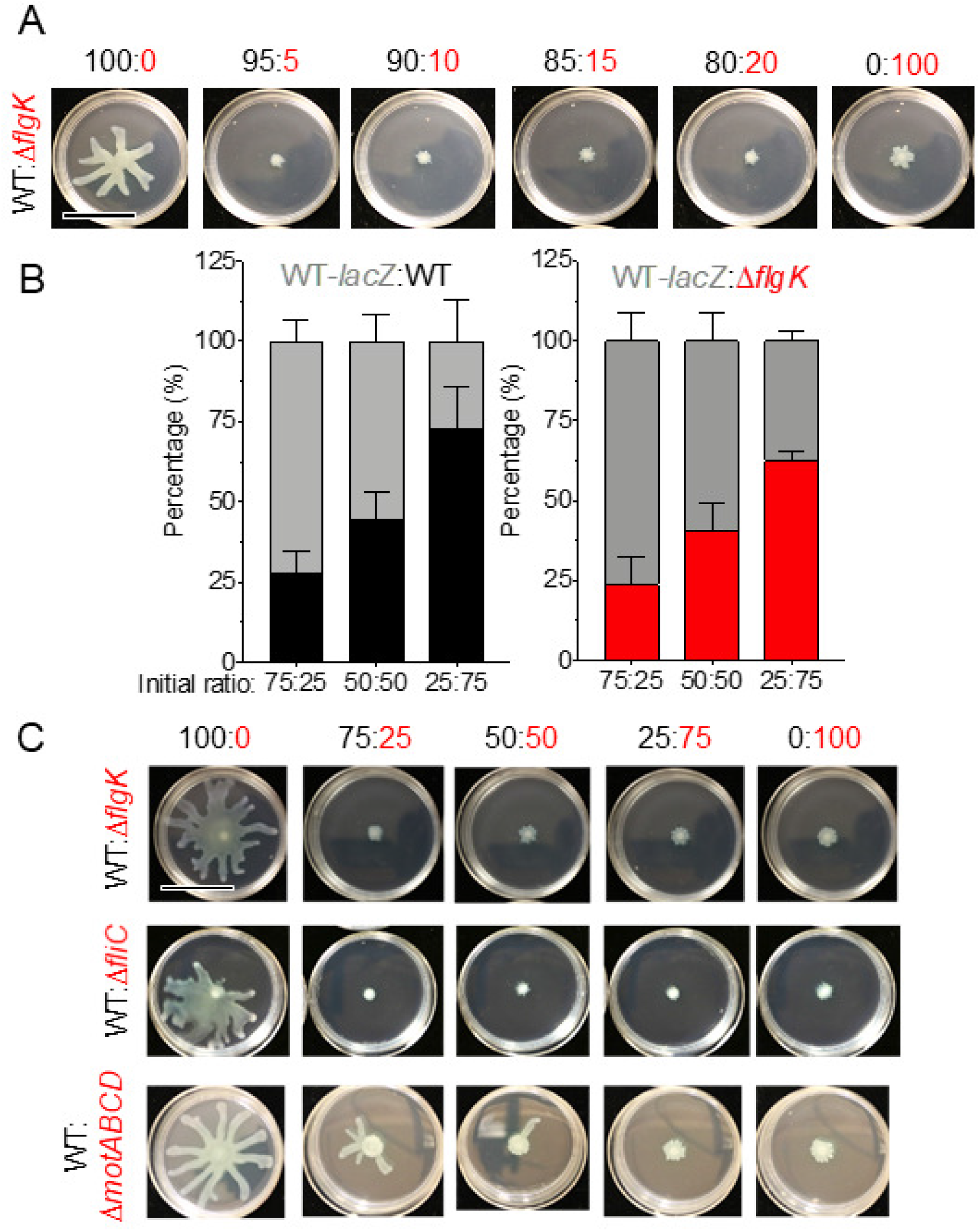
Motility heterogeneity represses swarming motility independent of competition. **A**. Representative images from swarm assay of wild-type (WT) *P. aeruginosa* strain PA14 (flagellated) mixed with Δ*flgK* (non-flagellated) at the indicated ratios on M8 agar. In this and all figure panels in the manuscript, WT and derivatives are labeled in black, while non-motile mutants are labeled in red. Data are representative of three individual experiments. **B**. Competition growth assay between WT tagged with a *lacZ* reporter (gray) and either an untagged WT strain (black) or Δ*flgK* mutant (red). Data show the average of three individual experiments. Error bars represent the standard deviation between the replicate values. Statistical analysis using one-way ANOVA showed no difference between input and output for each ratio analyzed. **C**. Representative images from swarm assay of WT (flagellated) mixed with Δ*flgK* (non-flagellated), Δ*fliC* (non-flagellated), or Δ*motABCD* (paralyzed flagellum) in the indicated ratios on M8 agar. Data are representative of three individual experiments. Scale bar: 30 mm

To determine if the lack of swarming in WT:Δ*flgK* co-cultures was due to faster growth of the non-flagellated cells compared to the WT, we competed the WT or the Δ*flgK* mutant against a WT strain tagged with a *lacZ* reporter at different ratios atop a 0.22 μm filter placed on 0.5% M8 agar. After 16 h, the colony was disrupted and colony forming units (CFUs) for each strain were enumerated on blue/white screening plates that contained X-Gal. At all ratios tested, neither untagged WT nor the Δ*flgK* mutant could outgrow the *lacZ*-expressing WT strain (**Fig. 1B**), indicating that the lack of swarming observed in WT:Δ*flgK* mixes was not due to increased growth of non-flagellated cells and thus overgrowth of the population. Rather, our data suggest that a subpopulation of cells incapable of flagellar motility inhibits motility of the larger flagellated WT population.

Using conditions in which the non-motile strain comprised 25%, 50%, or 75% of the population, we also found that, like the Δ*flgK* mutant, the Δ*fliC* mutant, another strain incapable of flagellar motility due to the lack of the flagellin protein used to make the flagellar filament, inhibited WT swarming at all ratios tested (**Fig. 1C**). Similarly, the Δ*motABCD* mutant, which produces a flagellum that is unable to rotate due to the absence of stators, repressed swarming of the WT; although, in comparison to Δ*flgK* and Δ*fliC*, the Δ*motABCD* mutant was slightly less effective at repressing WT swarming (**Fig. 1C**). Together, these data suggest that the absence of flagellar motility in a subpopulation of cells is sufficient to suppress WT swarming in co-culture.

### Functional T4P in flagellated cells are required for non-flagellated cells to repress swarming in co-culture

Several studies have shown that *P. aeruginosa* strains that are deficient in T4P tend to have a hyper-swarming phenotype (23-25), while hyperpiliated strains (those that overproduce elongated pili due to a defect in retraction) have inhibited swarming (15, 26). Additionally, *P. aeruginosa* cells have been shown to interact via their T4P to form close cell associations that facilitate directional swarming (15). Therefore, we determined if T4P are required by flagellated cells to facilitate the ability of Δ*flgK* to inhibit population-wide swarming motility. To do this, we analyzed the ability of Δ*flgK* to repress swarming when co-cultured with the following mutants: Δ*pilA*, which lacks T4P; Δ*pilMNOP*, which lacks the alignment complex required for T4P function (27) and cyclic-di-GMP signaling (28); and both Δ*pilT* and Δ*pilU*, hyperpiliated mutants that lack either of the ATPases that mediate T4P retraction (29). Strains lacking *pilA, pilMNOP, pilT*, and *pilU* were all defective for twitching motility (**Fig. S2** and **Fig. S3**). The swarming motility of strains lacking *pilA* or *pilMNOP* was no longer inhibited by the Δ*flgK* mutant in co-culture (**Fig. 2** and **Fig. S4**). Interestingly, despite its reported hyperpiliation and in contrast to previous reports (15), Δ*pilU* was capable of swarming and swarming motility was inhibited by the Δ*flgK* mutant in co-culture like the WT (**Fig. 2, Fig. S5**). However, the hyperpiliated Δ*pilT* mutant was incapable of swarming (**Fig. 2**), in agreement with previous models (15). Complementation of the Δ*pilU* and Δ*pilT* mutants restored twitching in both strains as well as swarming in Δ*pilT* (**Fig. S6**).

**Fig. 2.**
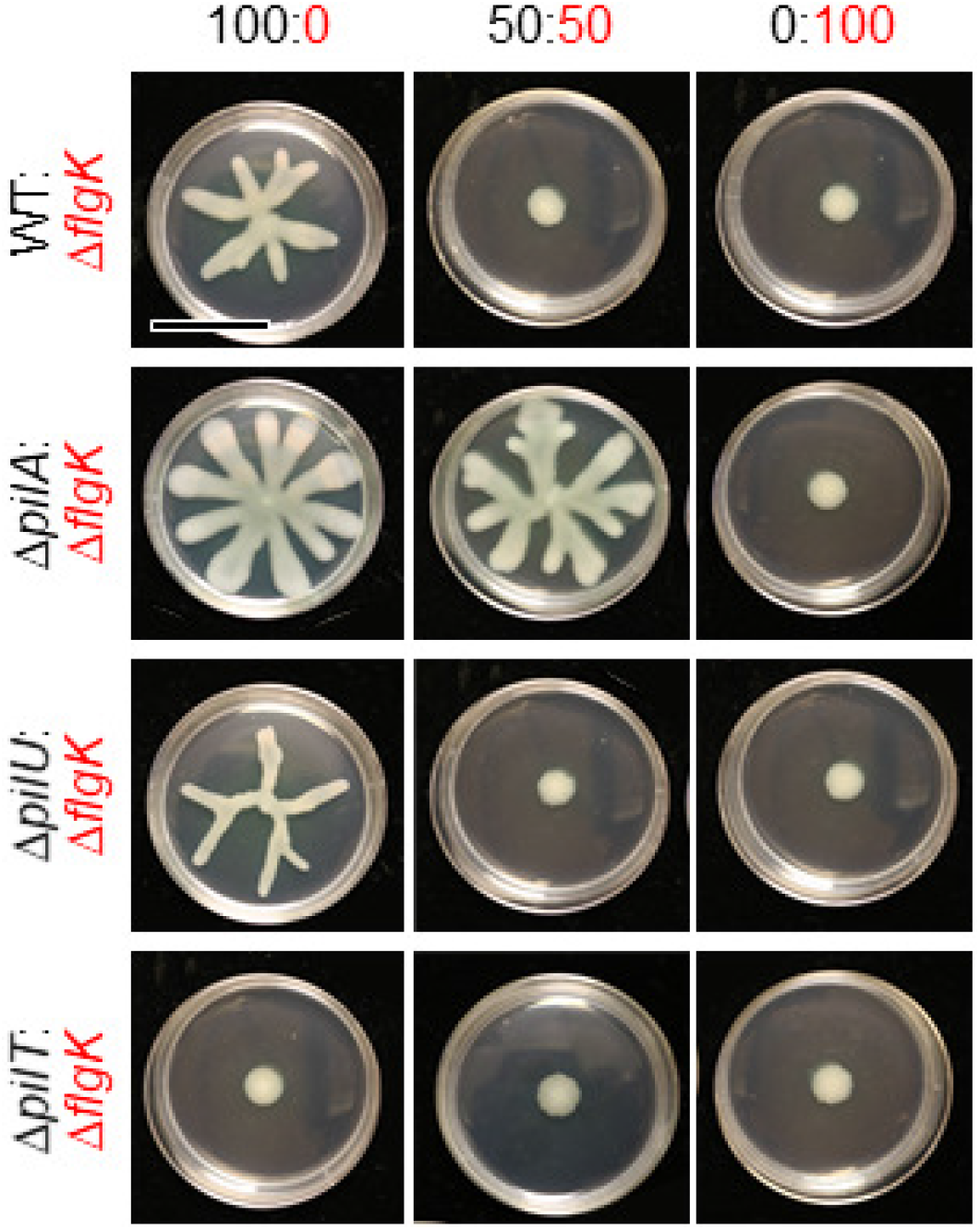
Flagellated *P. aeruginosa* requires functional T4P to be repressed by the non-motile subpopulation. Representative images from swarm assays of the non-flagellated Δ*flgK* mutant (functional T4P) mixed with either wild-type (WT) *P. aeruginosa* strain PA14 (functional T4P), the Δ*pilA* mutant (lacking T4P), the Δ*pilT* mutant (no T4P retraction), or the Δ*pilU* mutant (reduced T4P retraction) at the indicated ratios on M8 agar. Data are representative of three individual experiments. Scale bar: 30 mm

While PilT is the main retractile ATPase, PilU has an accessory role in retraction; the Δ*pilT* mutant completely lacks pilus retraction (30, 31), while the Δ*pilU* mutant has reduced retraction but still retains some function (31). This accessory role is supported by previous observations that the Δ*pilU* mutant retains sensitivity to infection by phage PO4, while the Δ*pilT* mutant became resistant to infection (26). We confirmed that the WT and Δ*pilU* mutant were sensitive to phage DMS3 infection and that the Δ*pilA* and Δ*pilT* mutants were resistant to infection under our experimental conditions (**Fig. S2B**). Additionally, while the Δ*pilU* mutant can form dense biofilms under static conditions, it is unable to form a biofilm under flow conditions likely due to Δ*pilU* T4P being more fragile than WT or even Δ*pilT* T4P, allowing for Δ*pilU* T4P to readily detach from the cell surface (26, 30). Overall, the data show that Δ*pilU* displays an intermediate phenotype in that it is hyperpiliated due to a reduction in T4P retraction that still results in inhibition of T4P-mediated twitching motility, but not swarming motility. Together, these data indicate that motile cells require T4P, and likely functional T4P, for the non-flagellated subpopulation to inhibit the swarming motility of the WT strain when grown in co-culture.

*P. aeruginosa* T4P participate in a cAMP-dependent signaling pathway (see **Fig. S7A** for pathway) that involves FimS, PilJ, CyaAB adenylate cyclases, and the cAMP-binding transcription factor Vfr (6). We found that this pathway was not required for the repression of WT swarming motility by the Δ*flgK* mutant (**Fig. S7B**) using published mutants lacking components of the cAMP signaling pathway (Δ*pilJ*, Δ*fimS*, Δ*cyaAB*, and Δ*vfr*) that retain expression of functional T4P as evidenced by their sensitivity to phage DMS3 (**Fig. S7C**). Taken together, the data show that flagellated WT cells require functional T4P, but not the cAMP-dependent surface-sensing pathway, in order for non-flagellated cells to be able to repress swarming in co-culture.’

### Non-flagellated cells require functional T4P to repress swarming motility of flagellated cells in co-culture

Consistent with published results (4, 32, 33), the Δ*flgK* mutant has functional T4P as evidenced by the formation of a large twitching motility zone using the agar-plastic interface assay (**Fig. S2A** and **Fig. S3**). While both the WT and Δ*flgK* formed twitch zones that were significantly larger than those formed by the T4P-null Δ*pilA* mutant, the Δ*flgK* twitch zone was ∼25% smaller than that formed by the WT (p<0.05; **Fig. S3A**). Using Δ*flgK* and Δ*flgK*Δ*pilA* strains, we found that *pilA*, which encodes the major pilin component of T4P, was required for the Δ*flgK* mutant to suppress WT swarming on 0.5% M8 agar (**Fig. 3A**). To assess whether Δ*flgK* cells needed functional pili to repress the WT, Δ*flgK* lacking either *pilU* (reduced T4P retraction) or *pilT* (no T4P retraction) were mixed with the WT. At all ratios tested, Δ*flgK*Δ*pilU* and Δ*flgK*Δ*pilT* were no longer able to repress WT swarming, like Δ*flgK*Δ*pilA*, indicating that the Δ*flgK* mutant required fully functional pili to repress population-wide swarming (**Fig. 3A, Fig. S8**). Complementation of *pilU* and *pilT* in Δ*flgK*Δ*pilU* and Δ*flgK*Δ*pilU*, respectively, was confirmed to restore twitching motility (**Fig. S9**); the effect of complementation on swarming motility was not assessed given that Δ*flgK* does not swarm. In a similar assay using 0.3% M63 agar, which supports both flagellum-mediated swimming and swarming, we found that inoculated spots of Δ*flgK* cells (**Fig. 3B, red dots**) decreased the local expansion of the motile WT population, while spots of the Δ*flgK*Δ*pilA* mutant (**Fig. 3B, purple dots**) did not. In contrast to the effects of Δ*flgK* cells on flagellar motility, the Δ*flgK* mutant did not alter T4P-dependent twitching motility of the WT in a 50:50 ratio when compared to the WT alone (**Fig. S3B**). Therefore, the data show that functional T4P are required for non-flagellated cells to inhibit swarming of WT cells in co-culture.

**Fig. 3.**
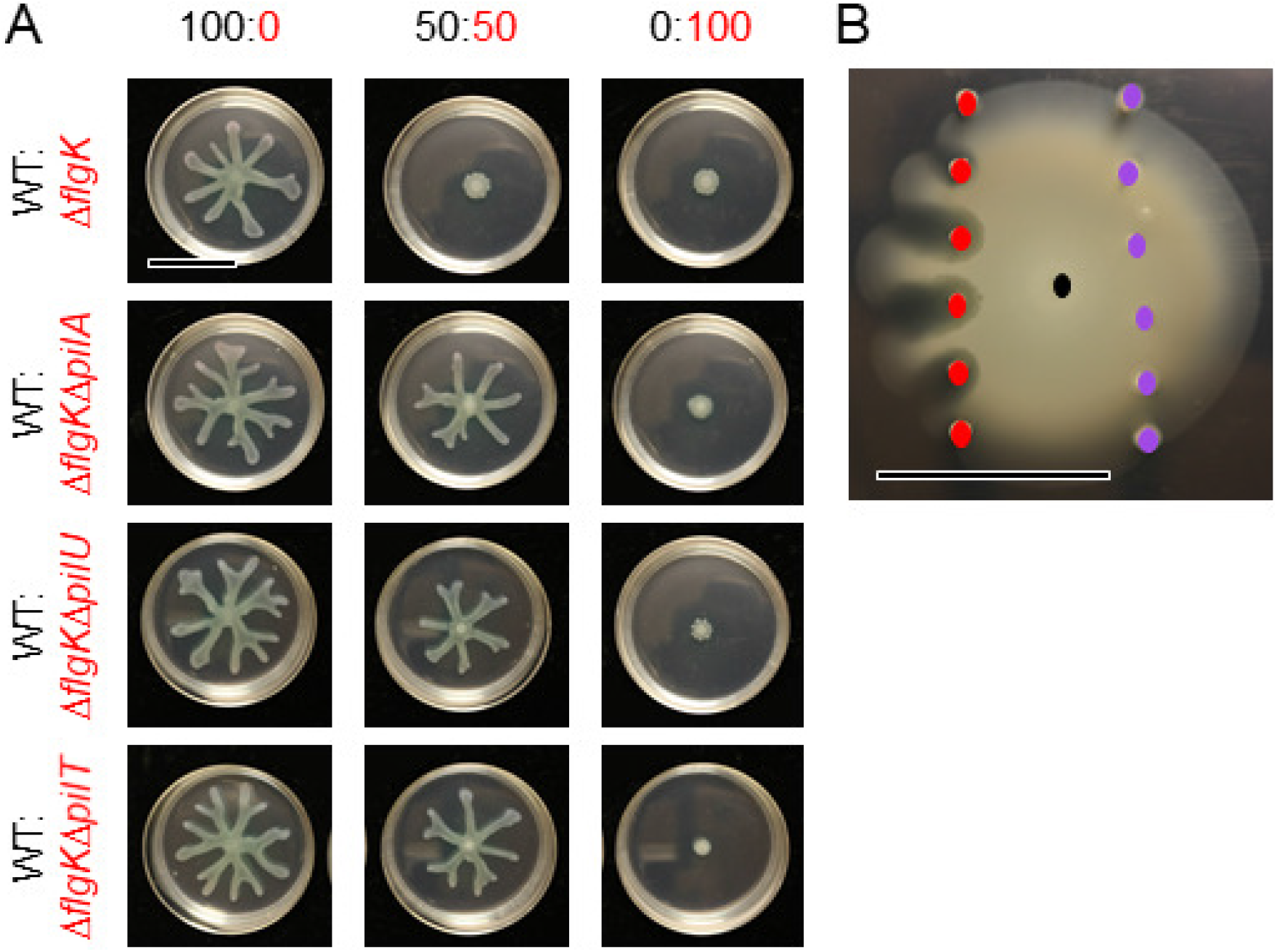
Non-flagellated *P. aeruginosa* requires T4P to repress flagellum-mediated motility of motile strains on soft agar. **A**. Representative images of swarm assays for wild-type (WT) *P. aeruginosa* strain PA14 (swarmer, Pel+, T4P+) mixed with Δ*flgK* (non-swarmer, Pel+, T4P+), Δ*flgK*Δ*pilA* (non-swarmer, Pel+, T4P-), Δ*flgK*Δ*pilU* (non-swarmer, Pel+, reduced T4P retraction), or Δ*flgK*Δ*pilT* (non-swarmer, Pel+, no T4P retraction) on M8 agar at the indicated ratios. WT is labeled in black while Δ*flgK* and derivatives are labeled in red. **B**. Representative image showing the interaction between WT (black dot), Δ*flgK* (red dots), and Δ*flgK*Δ*pilA* (purple dots) for flagellum-mediated swimming motility in soft agar (0.3% M63 agar) after 42 h incubation. The colored dots indicate the points of inoculation for the respective strains. Data are representative of three individual experiments. Scale bar: 30 mm

### Non-flagellated *P. aeruginosa* requires Pel matrix production to repress swarming motility in the flagellated population

We next explored whether Pel matrix production played a role in swarming repression by using a Δ*pelA* mutant, which is a mutant capable of robust swarming but lacks PelA-mediated deacetylase and hydrolase activities and, subsequently, secretion of properly modified Pel polysaccharide (34). We found that swarming by the Δ*pelA* mutant was repressed by Δ*flgK* in co-culture like the WT (**Fig. 4A**). In contrast, Δ*flgK*Δ*pelA* did not repress WT swarming (**Fig. 4A**). Flagellar mutants, such as Δ*fliC*, have been reported to overexpress Pel and Psl polysaccharides (35). Both Δ*flgK* and Δ*fliC*, which repress WT swarming (**Fig. 1**), had increased Congo Red binding, which is consistent with increased Pel or exopolysaccharide production (**Fig. 4B, Fig. S10**). Additionally, while Δ*flgK*Δ*pilA* (no T4P), Δ*flgK*Δ*pilU* (reduced T4P retraction), and Δ*flgK*Δ*pilT* (no T4P retraction) no longer repressed population-wide swarming motility (**Fig. 2A**), this does not appear to be due to a change in Pel production as all Δ*flgK* mutants displayed an increase in Congo Red binding (Pel production) (**Fig. 4B, Fig. S10**). Together, these data show that the non-flagellated strain needs to produce Pel matrix to repress swarming motility, but the flagellated strain does not.

**Fig. 4.**
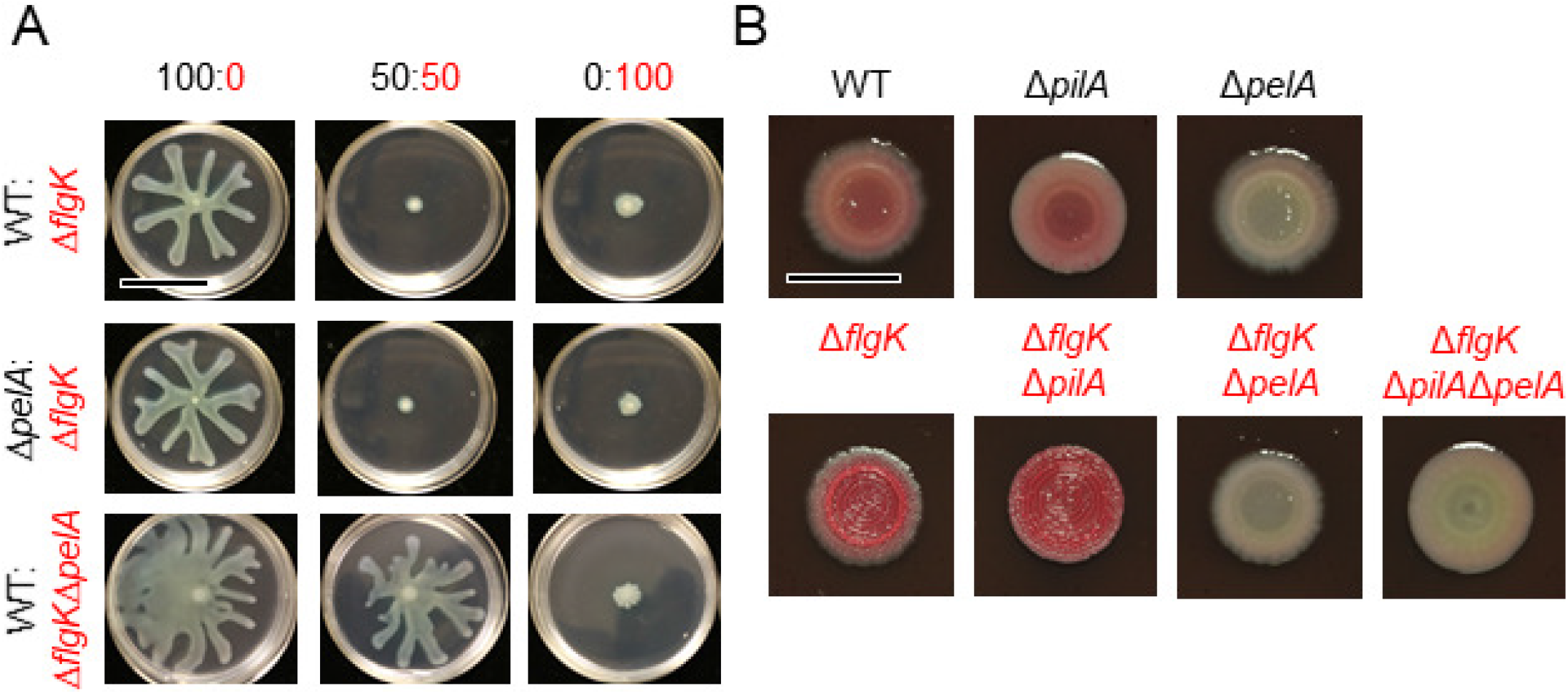
Pel matrix is only required by the non-flagellated *P. aeruginosa* subpopulation in order to repress overall swarming motility. **A**. Representative images from swarm assays of Δ*flgK* (non-flagellated and Pel+) mixed with wild-type (WT) *P. aeruginosa* strain PA14 (flagellated and Pel+) or Δ*pelA* (flagellated and Pel-) and of WT mixed with Δ*flgK*Δ*pelA* (non-flagellated and Pel-) at the indicated ratios on M8 agar. Scale bar: 30 mm **B**. Representative images of WT, Δ*pilA*, Δ*pelA*, Δ*flgK*, Δ*flgK*Δ*pilA*, Δ*flgK*Δ*pelA*, and Δ*flgK*Δ*pilA*Δ*pelA* grown on Congo Red plates to assess Pel production. WT and derivatives are labeled in black while Δ*flgK* and derivatives are labeled in red. Scale bar: 10 mm

## DISCUSSION

Based on the data presented here we propose a model (**Fig. 5**) whereby a subpopulation of cells defective in flagellar motility limit swarming motility of the entire population. Our analyses revealed that non-flagellated cells required T4P and Pel polysaccharide to inhibit the larger swarming population, and that flagellated cells required functional T4P but not the Vfr/cAMP signaling system to respond to the non-flagellated population. These findings in regards to T4P and Pel are consistent with the recruitment of cells to microcolonies during biofilm development and aggregate formation, which has been reported in numerous contexts (e.g. (4)). While cAMP was not required for repression of swarming, our previous findings showed that ethanol decreased swarming motility by increasing c-di-GMP (3). Future studies will investigate the role of c-di-GMP levels in the Δ*flgK*-mediated repression of WT swarming. Furthermore, given that Δ*flgK* requires retractable T4P and Pel to repress WT swarming, and overproduces Pel compared to the WT, future studies will test how varying the amount of Pel and TFP in WT and mutant populations contribute to repression of population-wide swarming.

**Fig. 5.**
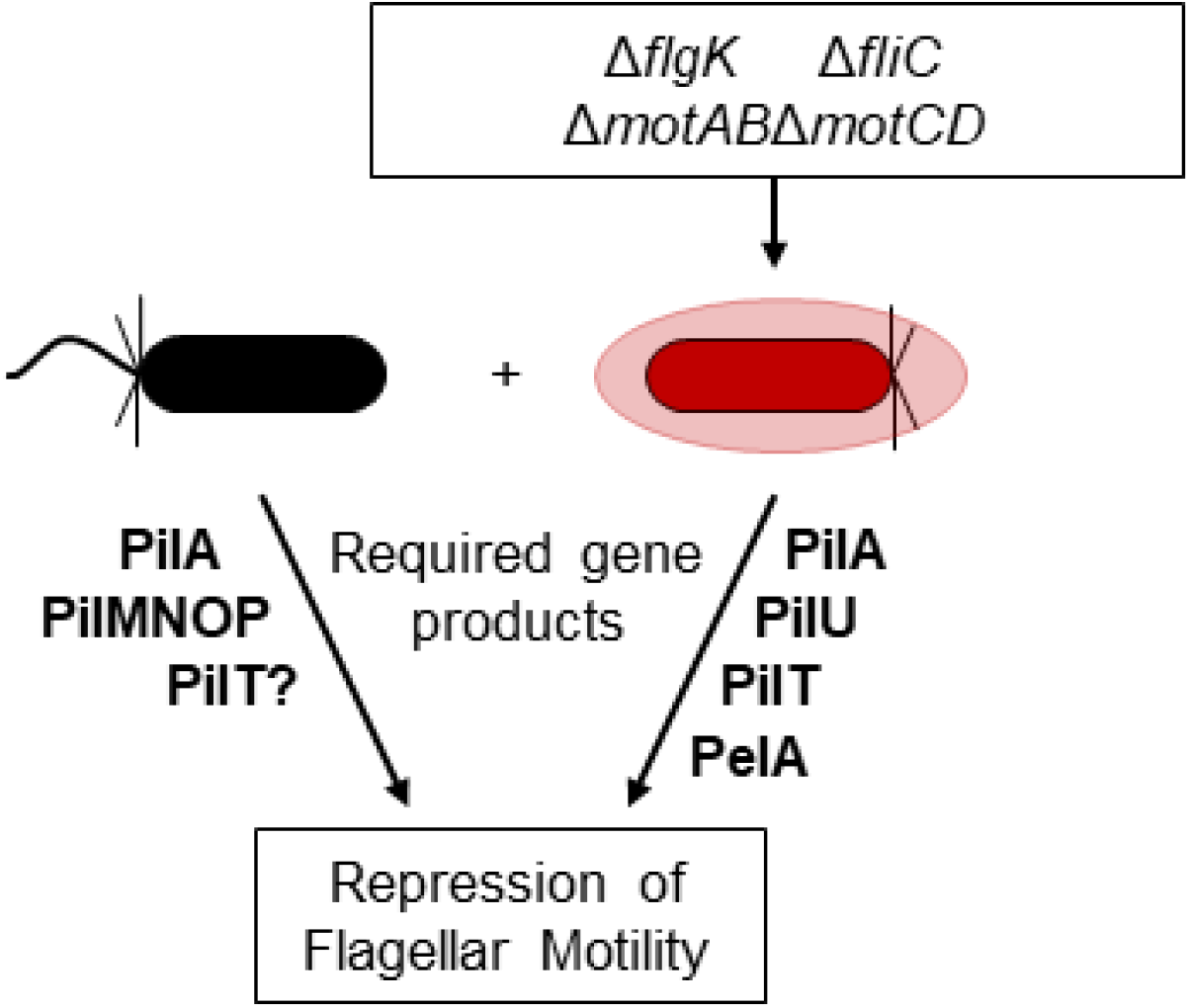
Summary model describing the genes and their products needed for repression of population-wide flagellum-mediated motility by non-flagellated cells in *P. aeruginosa* strain PA14. We show that a subpopulation of *P. aeruginosa* cells defective in flagellar motility (red) in co-culture with flagellated cells (black) impedes flagellum-mediated population-wide swarming motility in a manner that is dependent on retractable T4P (PilMNOP, PilA, PilU, PilT). We also show that Pel matrix (PelA), but only from the non-flagellated strain, is required. PilU, PilJ, FimS, CyaAB, and Vfr were not required for repression of swarming motility. These data may indicate ways that *P. aeruginosa* cells come together during biofilm formation.

The involvement of T4P in swarming cells was previously reported by Anyan et al. (15) who reported that T4P impact cell-cell interactions in swarms in ways that limit the movement of cells away from the population front. T4P mutants have been shown to have increased swarming motility, suggesting that the intercellular T4P interactions that we report between flagellated and non-flagellated genotypes may also be occurring in single-strain swarms in which all cells have the potential for flagellar motility (24, 25). Future studies will determine if T4P play direct roles in cell-cell interactions or indirect roles in structuring the community.

The WT and Δ*pelA* mutant responded similarly to non-flagellated cells while Δ*flgKpelA* was no longer able to repress motility suggesting that not all cells need to produce exopolysaccharide matrix to repress swarming. Our findings that Pel production is only necessary in the non-flagellated subpopulation is supported by published data showing that ethanol not only decreases *P. aeruginosa* flagellar motility (3), but increases Pel production (36). Pel has been suggested to repress motility via the steric hinderance of flagella (3, 37), but further studies are needed to test this hypothesis. The involvement of Pel in the interaction between flagellated and non-flagellated cells is particularly interesting in light of recent work by Whitfield et al. (38) which found that the Pel biosynthetic locus is widespread across Gram-positive and Gram-negative bacteria. Future studies will determine if Pel is specifically detected by a T4P-dependent mechanism or if other matrix materials play a similar role.

Boyle et al. (16) previously reported that Δ*flgK* was capable of repressing WT swarming when Δ*flgK* was added 5:1 or at 83% of the population, while we show that Δ*flgK* at as low as 5% of the population represses WT swarming. Our previously published findings (3) also showed that while strain PA14 was capable of swarming, an average of 38% of flagellated cells were motile at any given time. Furthermore, we showed that ethanol strongly repressed *P. aeruginosa* swarming motility by reducing the motile fraction of the population from 38% to 22% (3). Overall, the data show that motility of individual flagellated WT cells is heterogeneous within a population, but the addition of non-flagellated cells represses population-wide motility. These findings suggest that the number of non-flagellated cells necessary to repress WT motility may vary due to experimental conditions such as medium type and composition.

In the studies here, we seed a low percentage of non-swarming cells into a swarming population, which is reminiscent of the situations where there are genetically heterogeneous populations observed in the CF airway or other chronic infections (19, 39, 40) as well as during normal environmental growth (40, 41). Additionally, clinical *P. aeruginosa* isolates from the CF lung have diverse motility phenotypes (19). This diversity in motility is even observed within a single patient (19). We hypothesize that such a mechanism promotes interspecies and intraspecies interactions. The strong implication of these findings is that in a mixed population with a subpopulation of non-flagellated cells, the entire population can be brought to a halt. Similarly, within a population in which a significant number of cells have stopped swimming or swarming, the further addition of even just a few non-flagellated cells can suppress motility for the entire population (i.e., see the ethanol studies mentioned above). As *P. aeruginosa* populations inherently have some number of non-motile or inactive cells (3), future studies are required to determine how strain background or mutant genotype affects phenotypic heterogeneity among isogenic cells.

## Materials and Methods

### Strains and media

Strains used in this study are listed in **Table 1**. *P. aeruginosa* strain PA14 and derivatives were routinely cultured on lysogeny broth (LB) solidified with 1.5% agar or in LB broth at 37°C with shaking. For *P. aeruginosa* phenotypic assays, either M63 [22 mM KH_2_PO_4_, 40 mM K_2_HPO_4_, and 15 mM (NH_4_)_2_SO_4_] or M8 (42 mM Na_2_PO_4_, 22 mM KH_2_PO_4_, and 8.5 mM NaCl) minimal salts medium supplemented with MgSO_4_ (1 mM), glucose (0.2%), and casamino acids (CAA; 0.5%) were used as indicated. Complemented strains containing genes regulated by an arabinose-inducible promoter (P_*BAD*_) were grown in the presence of 0.25% arabinose.

**Table 1.**
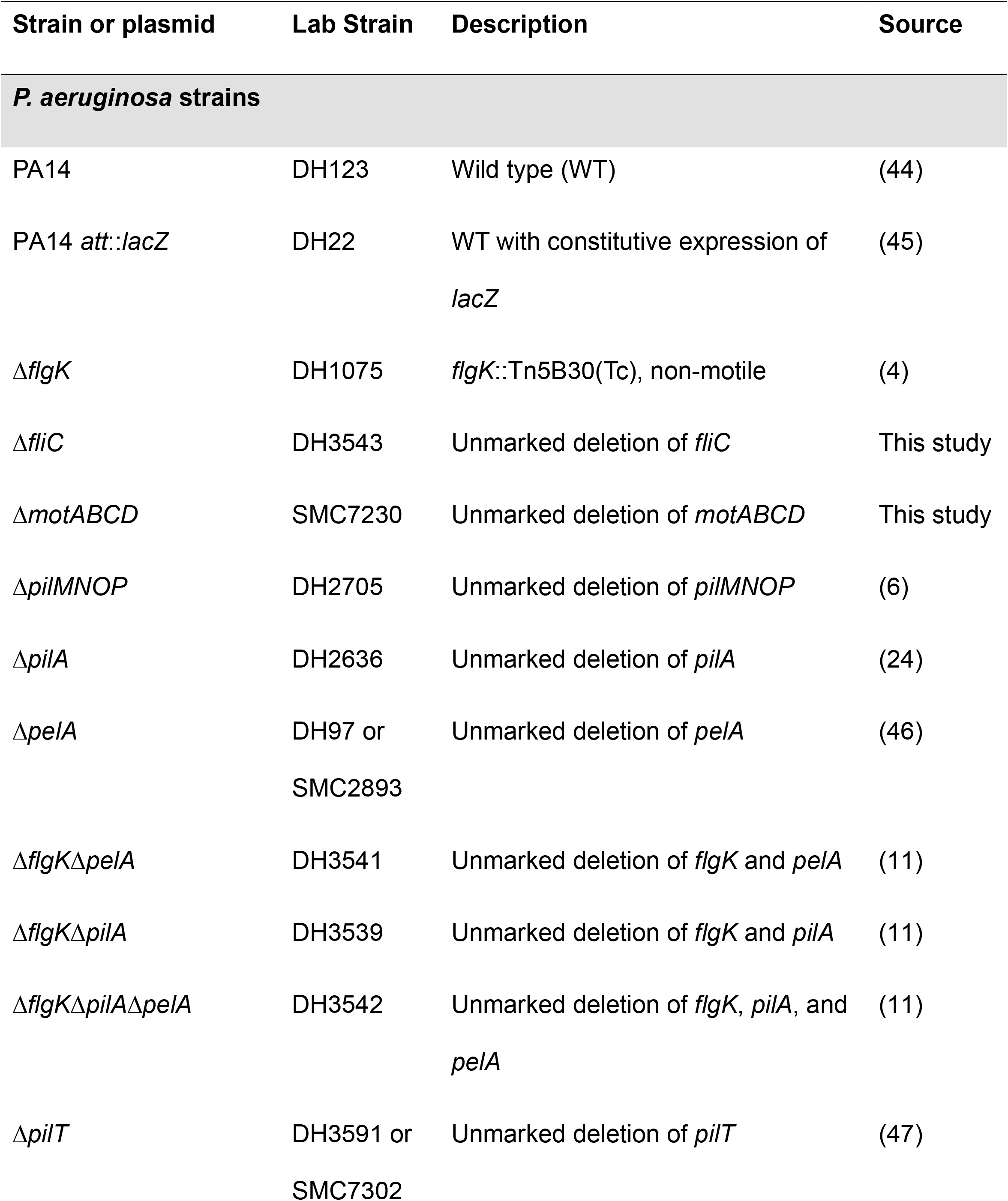
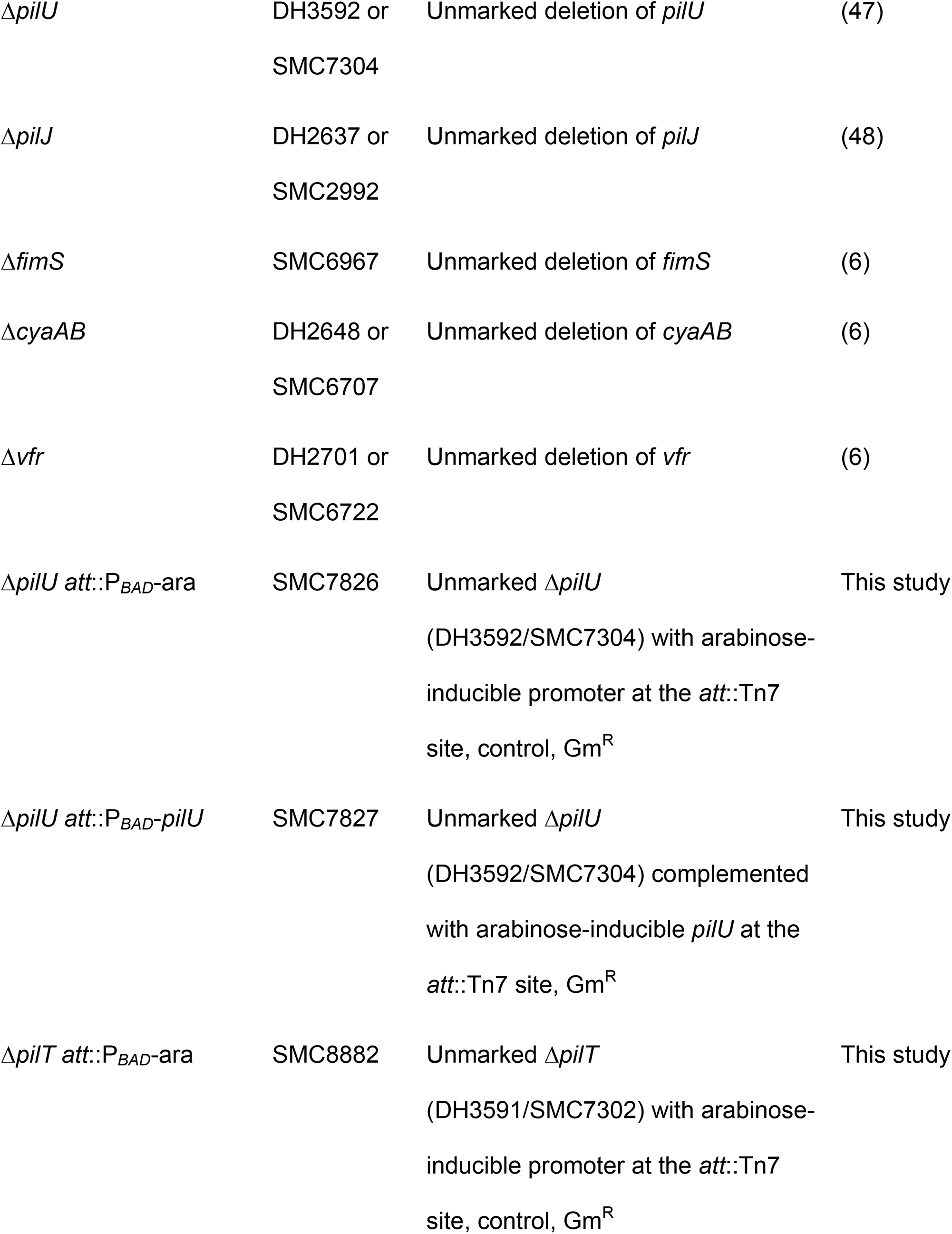
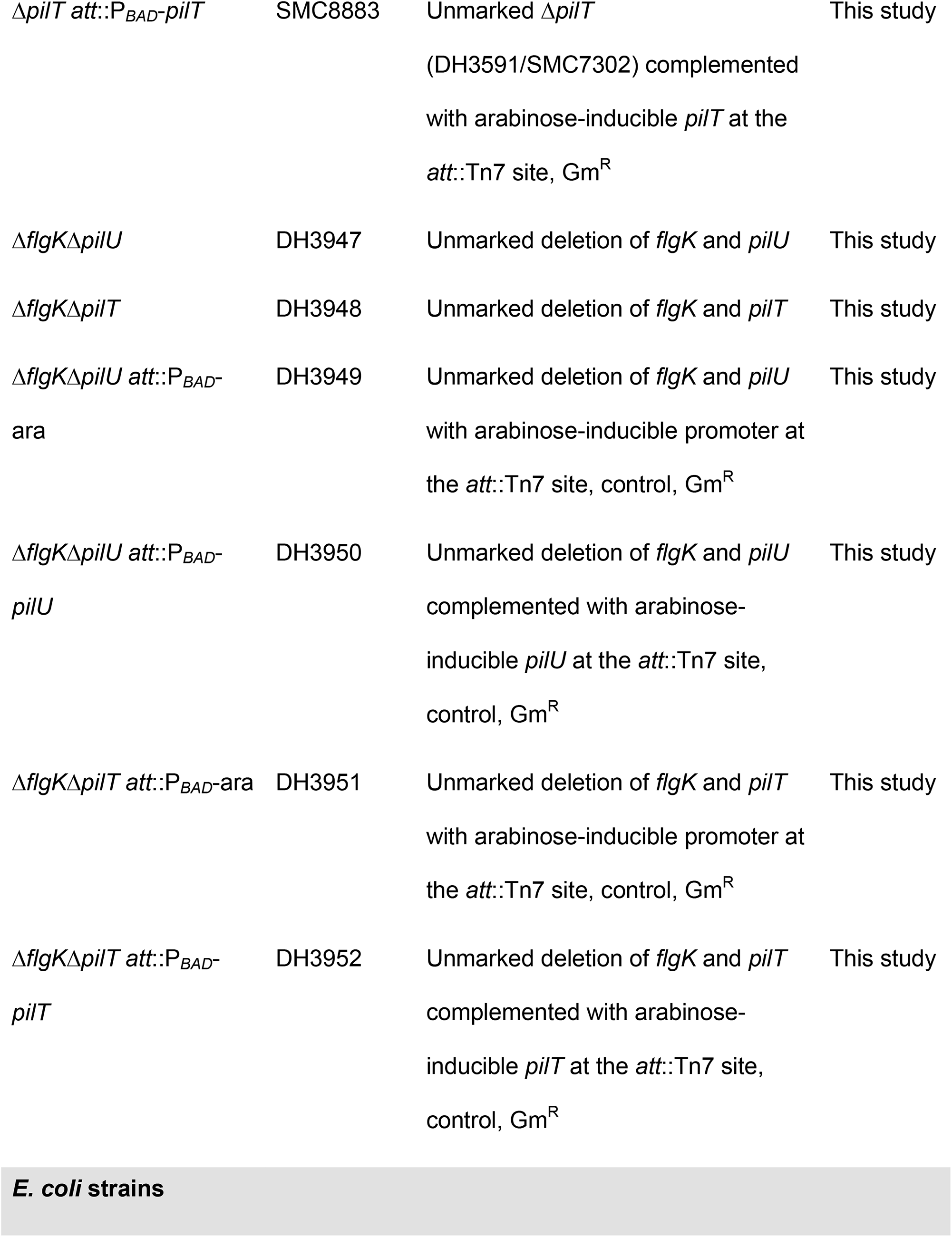
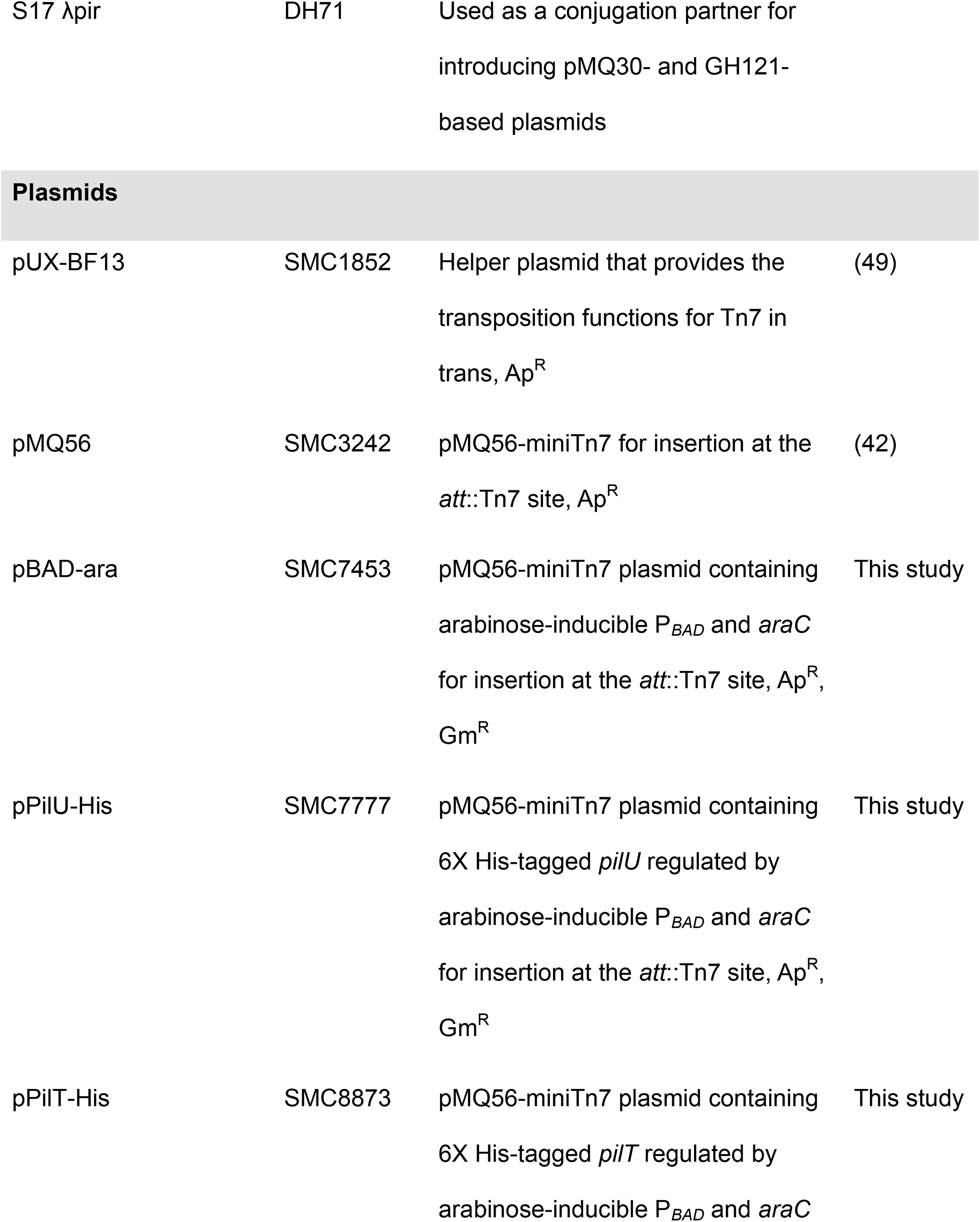
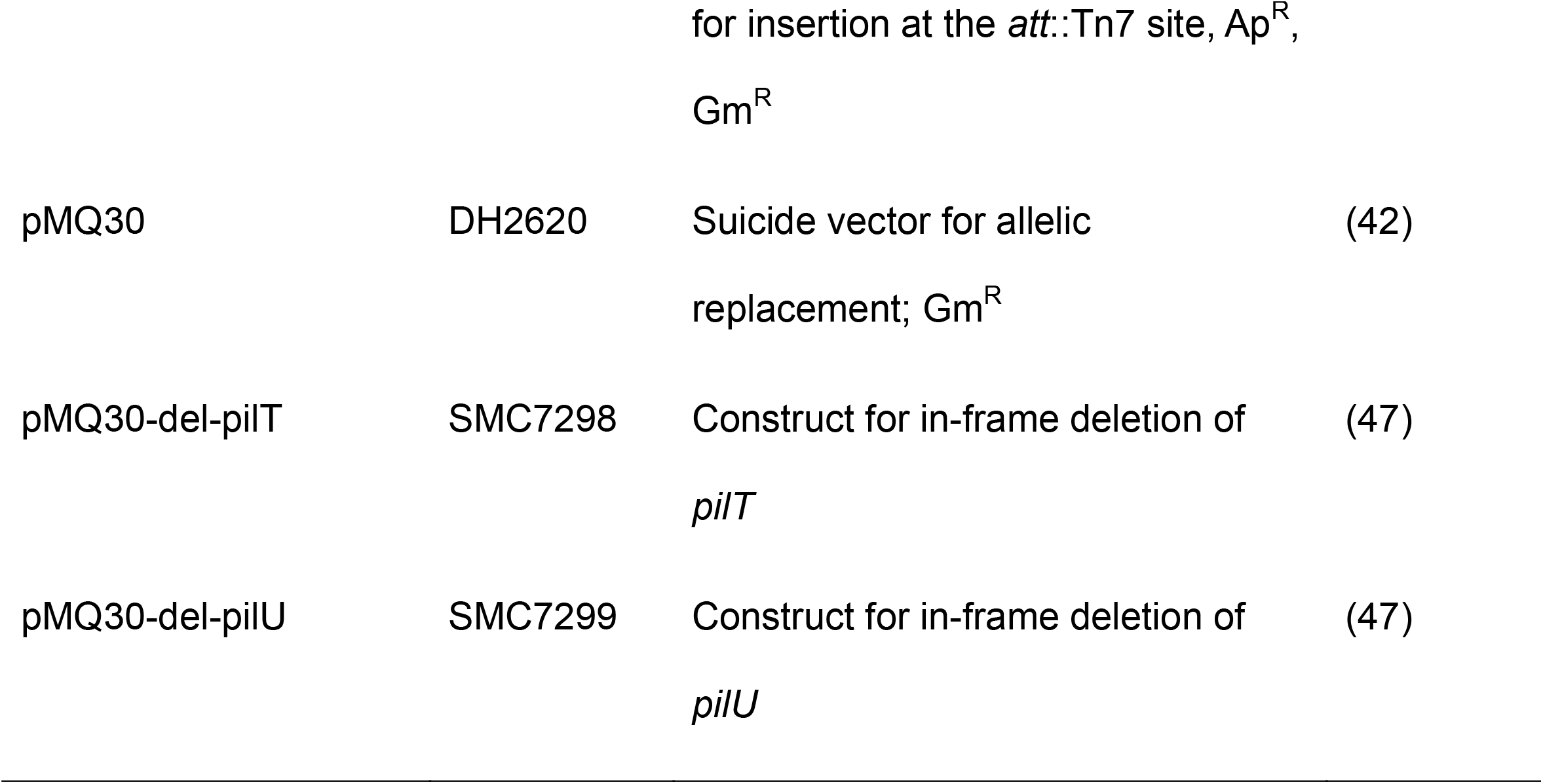
Strains and plasmids used in this study.

### Construction and complementation of mutant strains

Plasmids used in this study are listed in **Table 1**. Plasmid constructs for making in-frame deletions and arabinose-inducible complementation strains were constructed using a *Saccharomyces cerevisiae* recombination technique described previously (42) or Gibson Assembly. All plasmids were sequenced at the Molecular Biology Core at the Geisel School of Medicine at Dartmouth. Plasmids were introduced into *P. aeruginosa* by conjugation via S17/lambda pir *E. coli*. Merodiploids were selected by drug resistance and double recombinants were obtained using sucrose counter-selection and genotype screening by PCR.

### Swarming motility assays

Swarm assays were performed as previously described (43). Briefly, 9 ml of M8 medium with 0.5% agar (swarm agar) was poured into 60 × 15 mm plates and allowed to dry at room temperature for 3 h prior to inoculation. The indicated strains were grown for 16 h then each was washed in 1X PBS and then normalized to an OD_600_ of 1. Indicated isolates were then mixed at the indicated ratios in a final volume of 100 µl. Each plate was inoculated with 0.5 µl of the liquid culture mixture and the plates incubated face-up at 37°C in stacks of no more than four plates for up to 22 h. Each mixture was inoculated in three to four replicates and was assessed on at least three separate days. Images were captured using a Canon EOS Rebel T6i camera and images assessed for swarm repression.

### Swimming motility assays

Swim assays were performed as previously described (43). Briefly, M63 medium solidified with 0.3% agar (soft agar) was poured into 100 × 15 mm petri plates and allowed to dry at room temperature (∼25°C) for 3 h prior to inoculation. Sterile toothpicks or P20 pipette tips were used to inoculate bacterial mixes into the center of the agar without touching the bottom of the plate; liquid cultures grown for 16 h were used as inoculum. The indicated strains were normalized to an OD_600_ of 1 then were mixed at the indicated ratios in a final volume of 100 µl. No more than four bacterial mixtures were assayed per plate. Plates were incubated upright at 37°C in stacks of no more than four plates per stack for 18 to 20 h.

### Twitching motility assays

T-broth [1% tryptone (w/v), 0.5% NaCl (w/v)] medium solidified with 1.5% agar was poured into petri plates and allowed to dry at room temperature (∼25°C) for 3 h. Overnight (16 h) cultures were washed once in 1X PBS and then normalized to an OD_600_ of 1. Plates were inoculated by dipping a sterile toothpick or P20 pipette tip into the washed and normalized cultures and then inserting the toothpick into the agar until it touched the bottom of the plate. Plates were then incubated at 37°C for 46 h after which time the agar was removed and the plate incubated in 0.1% crystal violet for 10 min. Plates were then washed three times in water and dried at room temperature. Images of the dried, stained twitch area were taken and twitch diameter was measured.

### Competition experiments

Competition assays were performed to determine relative growth of selected *P. aeruginosa* strains. Strains were grown for 16 h and then 1 ml of culture was pelleted at 15,682 *x g* for 2 min, and washed once in 1 ml 1X PBS followed by resuspension in 1ml 1X PBS. The OD_600_ of each culture was normalized to 1. The strains to be competed were mixed at the indicated ratios in a final volume of 100 µL and then 0.5 µL of the combined suspension was spotted onto a 0.22 µm polycarbonate filter (Millipore) placed on the surface of a swarm plate in triplicate. Plates were incubated at 37°C. Filters were then transferred into a 1.5 ml tube and the filter-associated cells were resuspended by adding 1 ml 1X PBS + 0.05% Triton X-100 detergent and agitating the tubes at high speed for 2 minutes using a Genie Disruptor (Zymo). This suspension as well as the inoculum were diluted, spread on LB plates supplemented with 150 µg/ml 5-bromo-4-chloro-3-indolyl-β-D-galactopyranoside (X-Gal) using glass beads, and incubated at 30°C until blue colonies were visible (∼24 h). The number of blue and white colonies per plate were counted and recorded to determine the relative abundance of each strain.

### Phage susceptibility assays

Phage susceptibility was analyzed by the cross-streak method or by spotting phage directly onto a lawn of *P. aeruginosa*. For the cross-streak method, *P. aeruginosa* strains were grown for 16 h in LB before diluting 1:100 in LB and then growing to an OD_600_ of 0.5-0.7. Then, 4 µl of phage DMS3 vir strain was spotted on the side of an LB agar plate and then dragged across the agar surface in a straight line before allowing to absorb into the agar. Once absorbed, 4 µl of each *P. aeruginosa* strain was spotted at the top of the agar and dragged downward in a straight line through the phage line. Plates were incubated at 37°C for 16 h.

Alternatively, 1% M8 agar plates (60 x15 mm) containing 0.2% glucose (v/v), 0.5% casamino acids (v/v), and 1 mM MgSO_4_ were prepared and cooled to room temperature. 50 µl of *P. aeruginosa* 16 h cultures were added to 1 ml of 0.5% warm top agar (M8 medium and supplements). The mixture was gently mixed and quickly poured onto M8 agar plates. Plates were swirled to ensure even spreading of top agar. Once cooled, 2 µl of phage DMS3 vir strain were spotted onto the center of the plate and allowed to dry before incubating plates at 37°C for 16 h.

### Congo red assay

Cultures were grown in 5 ml LB for 16 h at 37°C with rolling. Culture aliquots were washed once and resuspended in sterile deionized water. Washed cells were spotted (3 µl) onto Congo Red plates (1% tryptone [w/v], 1.5% agar, 40 µg/ml Congo Red, 20 µg/ml Coomassie Blue). Plates were grown for 16 h at 37°C and then moved to room temperature for 3 days to allow for color development.

### Statistical analysis

One-way ANOVA with multiple comparisons was performed pairwise between all isolates and mixtures using the GraphPad Prism 6 software (GraphPad, La Jolla, CA).

## Acknowledgements

This work was supported by CFF HOGAN19G0 to DAH, R37AI83256 and R01AI43730 to GAO, and CFF VERMIL21F0 to DMV. This work was also supported by P30DK117469 for the Applied Bioinformatics and Biostatistics Core. Sequencing services and specialized equipment was provided by the Genomics and Molecular Biology Shared Resource Core at Dartmouth supported by NCI Cancer Center Support Grant 5P30CA023108-37. Equipment used was supported by the NIH IDeA award to Dartmouth BioMT P20GM113132. We thank Amy Baker for constructing the Δ*fliC* mutant and Christine Toutain for construction of the *motABCD* deletion strain.

