## Supplemental Material for "Non-motile subpopulations of *Pseudomonas aeruginosa* repress flagellar motility in motile cells through a type IV pili- and Pel-dependent mechanism"

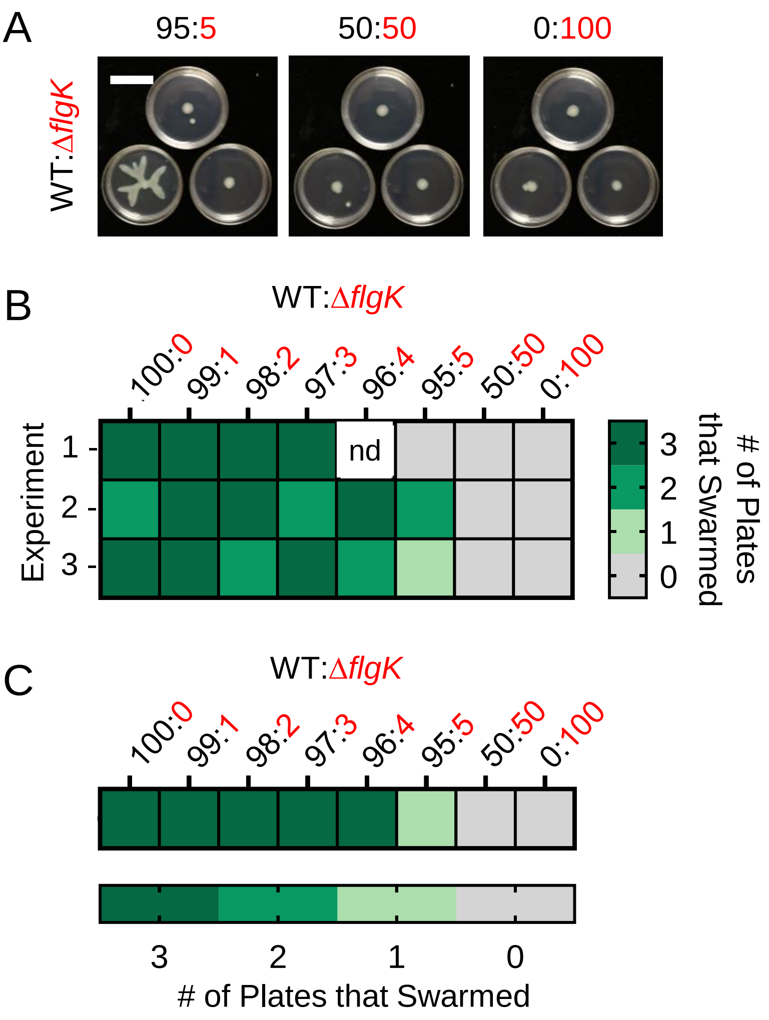


**Fig. S1. Establishing the lower limit of Δ*flgK* cells that is sufficient to repress motility A.** Representative images from swarm assays of wild-type *P. aeruginosa* strain PA14 (WT) mixed with Δ*flgK* (non-swarmer) at 95:5, 50:50, and all Δ*flgK* mutant. Images show three technical replicates from a single experiment. Consistent with published observations, there can be heterogeneity in swarm phenotypes despite best efforts for uniform assay conditions. Scale bar: 30 mm. **B.** Heatmap showing the number of plates (technical replicates) that swarmed at each ratio in three individual experiments (1-3). **C.** Heatmap showing the average number of plates (technical replicates) that swarmed at each ratio using data in **B**. nd, not determined.

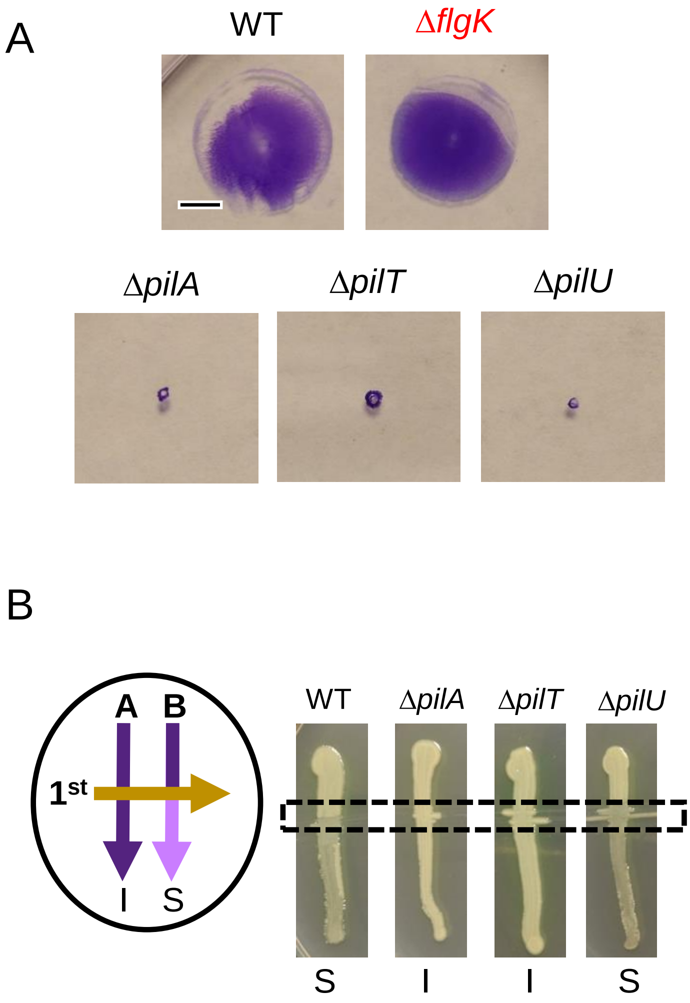


**Fig. S2. A.** Representative images from twitch assays of wild-type (WT) *P. aeruginosa* strain PA14, Δ*flgK, ΔpilA, ΔpilT, and ΔpilU* on T-broth agar after 46 h incubation at 37$^{o}$C. **B.** Cross-streak assay set up. On an LB plate, the phage lysate is first streaked horizontally (orange arrow) followed by vertical streaks of each bacterial strain. ”S” denotes strains that are sensitive to phage uptake and lysis (lighter cell density, light purple) and “I” denotes strains that are insensitive to phage uptake and lysis (heavier cell density, dark purple). Representative images of the cross-streak assay of WT, Δ*pilA*, Δ*pilT*, and Δ*pilU* showing sensitivity and insensitivity to phage DMS3. Dashed line indicates phage line. WT and derivatives are labeled in black while Δ*flgK* is labeled in red. Data are representative of three individual experiments showing similar results. Scale bar: 5 mm

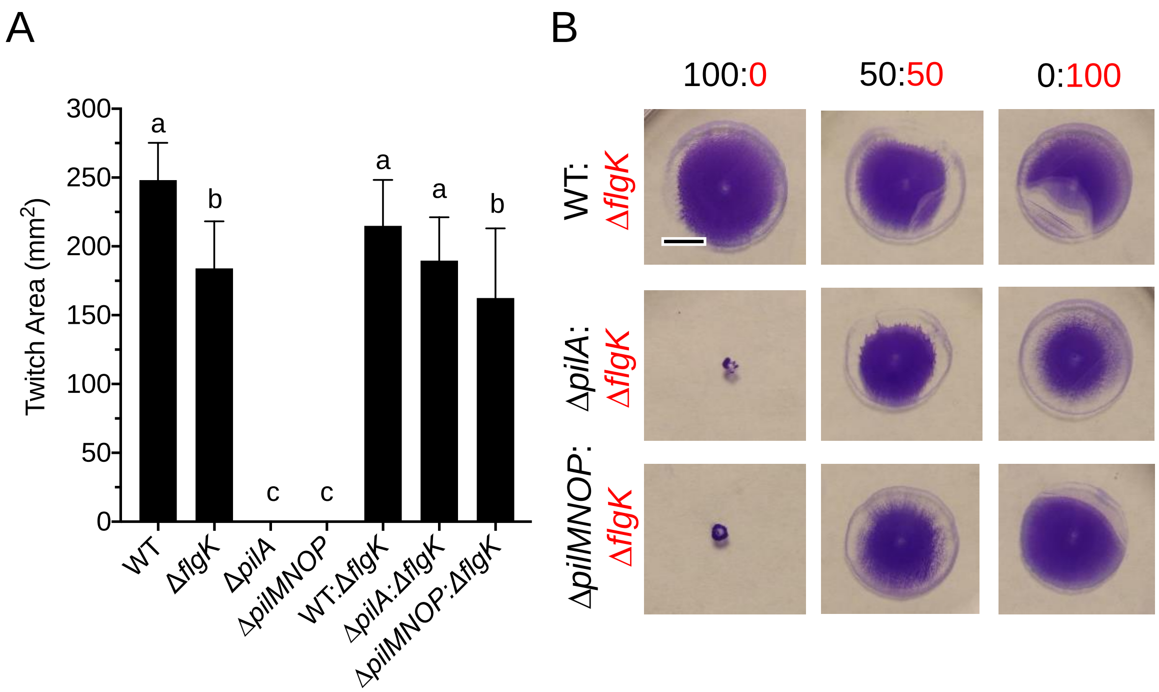


**Fig. S3. Motility heterogeneity does not impact T4P-mediated twitching motility.** **A.** Quantified results of the twitch assay of individual and mixed samples of the following strains: wild-type (WT) *P. aeruginosa* strain PA14, Δ*flgK*, Δ*pilA*, and Δ*pilMNOP*. Mixed samples were done at a 50:50 ratio. Same letters indicate samples that are not significantly different, while different letters indicate significant differences ($p$<0.05), as determined by One-way ANOVA with multiple comparisons. **B.** Representative images from twitch assays of WT (flagellated and T4P+), Δ*pilA* (flagellated and T4P-), and Δ*pilMNOP* (flagellated and T4P-) mixed with Δ*flgK* (non-flagellated and T4P+) at the indicated ratios on T-broth agar. WT and derivatives are labeled in black while Δ*flgK* is labeled in red. Data are **A.** the average or **B.** representative of three individual experiments. Scale bar: 5 mm

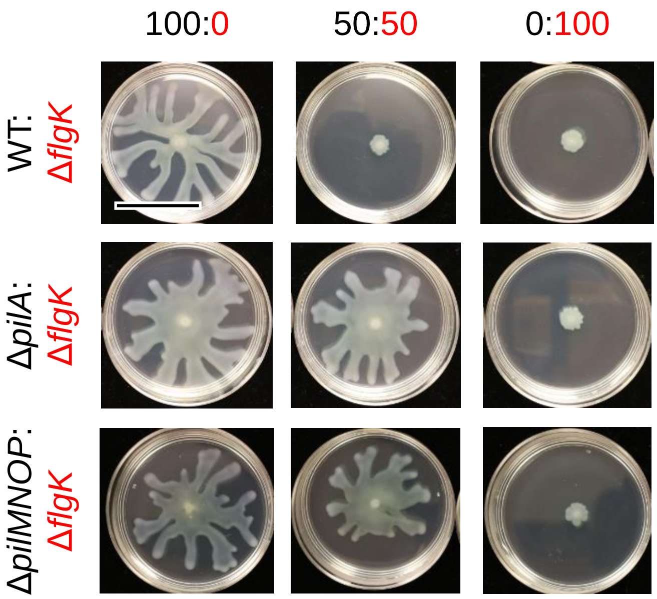


**Fig. S4. Flagellated *P. aeruginosa* requires both *pilA* and *pilMNOP* for non-flagellated Δ*flgK* to repress swarming.** Representative images from swarm assays of wild-type (WT) *P. aeruginosa* strain PA14 (flagellated and T4P+), Δ*pilA* (flagellated and T4P-), or Δ*pilMNOP* (flagellated and T4P-) mixed with Δ*flgK* (non-flagellated and T4P+) at ratios indicated. WT and derivatives are labeled in black while Δ*flgK* is labeled in red. Data are representative of three individual experiments showing similar results. Scale bar: 30 mm

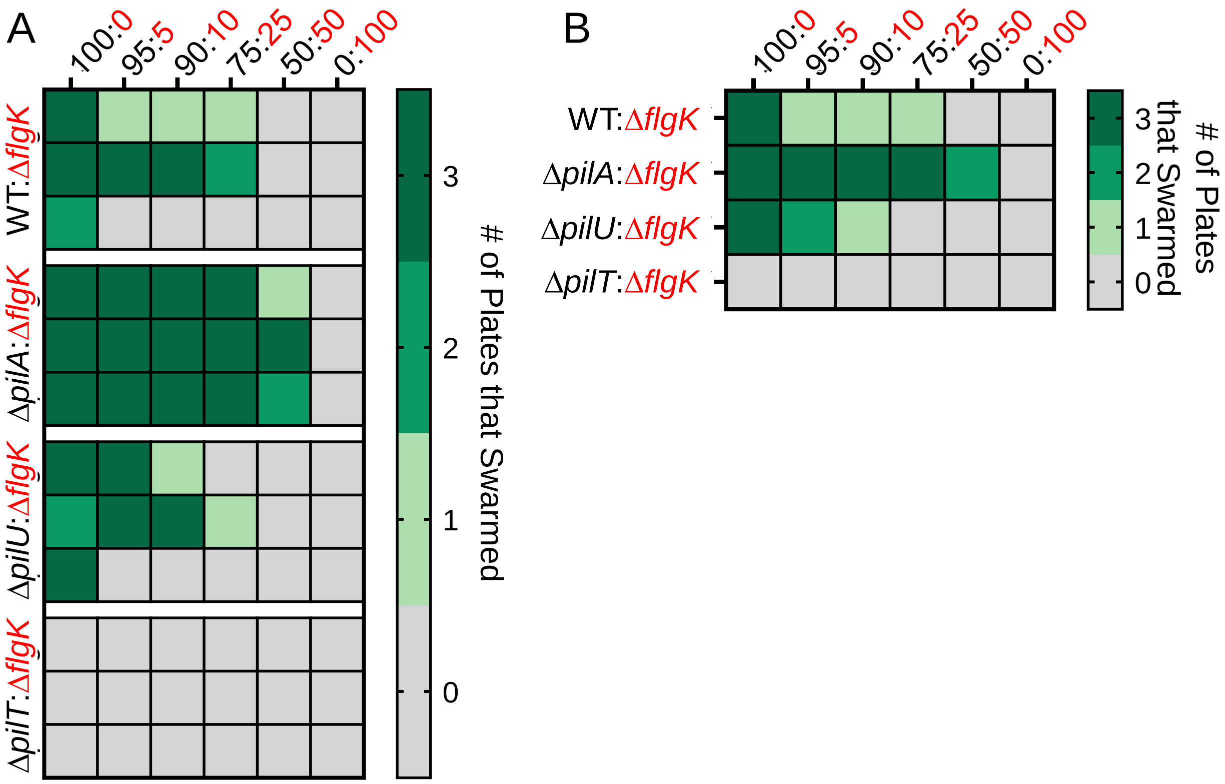


**Fig. S5. The hyperpiliated Δ*pilU* mutant with reduced T4P retraction swarms and is repressed by Δ*flgK*, but the hyperpiliated Δ*pilT* mutant with no T4P retraction is unable to swarm. A.** Heatmap showing the number of plates (technical replicates) that swarmed at each ratio for three individual experiments represented by three different rows. **B.** Heatmap showing the average number of plates (technical replicates) that swarmed at each ratio using data in **A**. WT and its derivatives are labeled in black while Δ*flgK* is labeled in red.

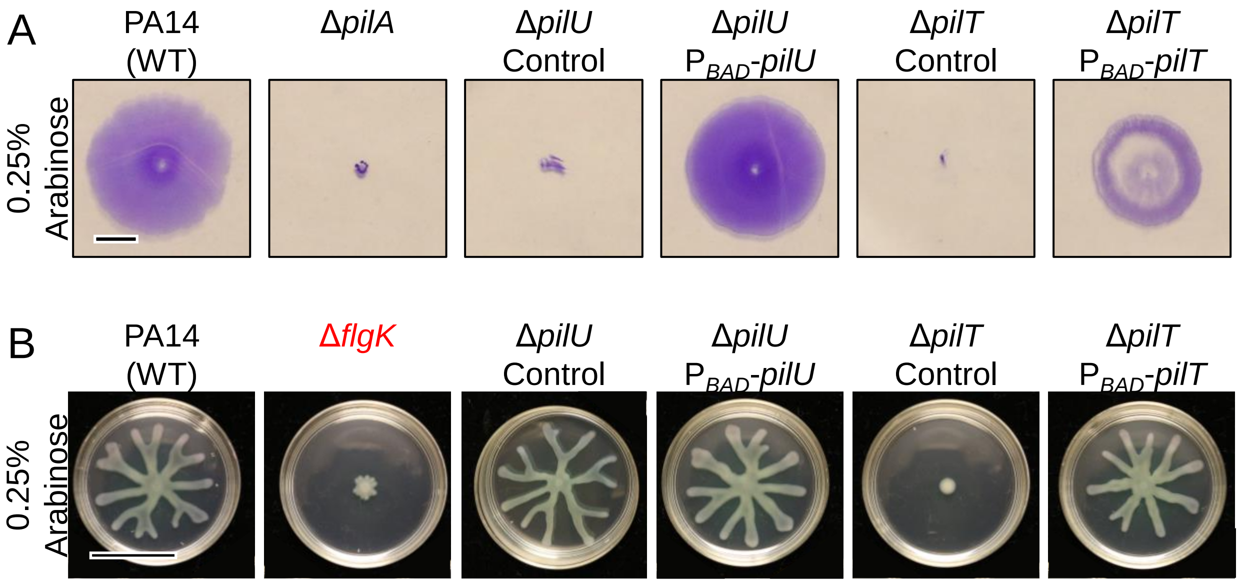


**Fig. S6. Complementation of Δ*pilU* and Δ*pilT***. Representative images from **A.** twitch assays and **B.** swarm assays of wild-type (WT) *P. aeruginosa* strain PA14, Δ*pilA,* Δ*pilU,* Δ*pilU* P$_{BAD}$-*pilU* (complemented), Δ*pilT*, and Δ*pilT* P$_{BAD}$-*pilT* on T-broth agar containing 0.25% arabinose after 46 h incubation at 37$^{o}$C. Data are representative of three individual experiments. **A.** Scale bar: 5 mm **B.** Scale bar: 30 mm

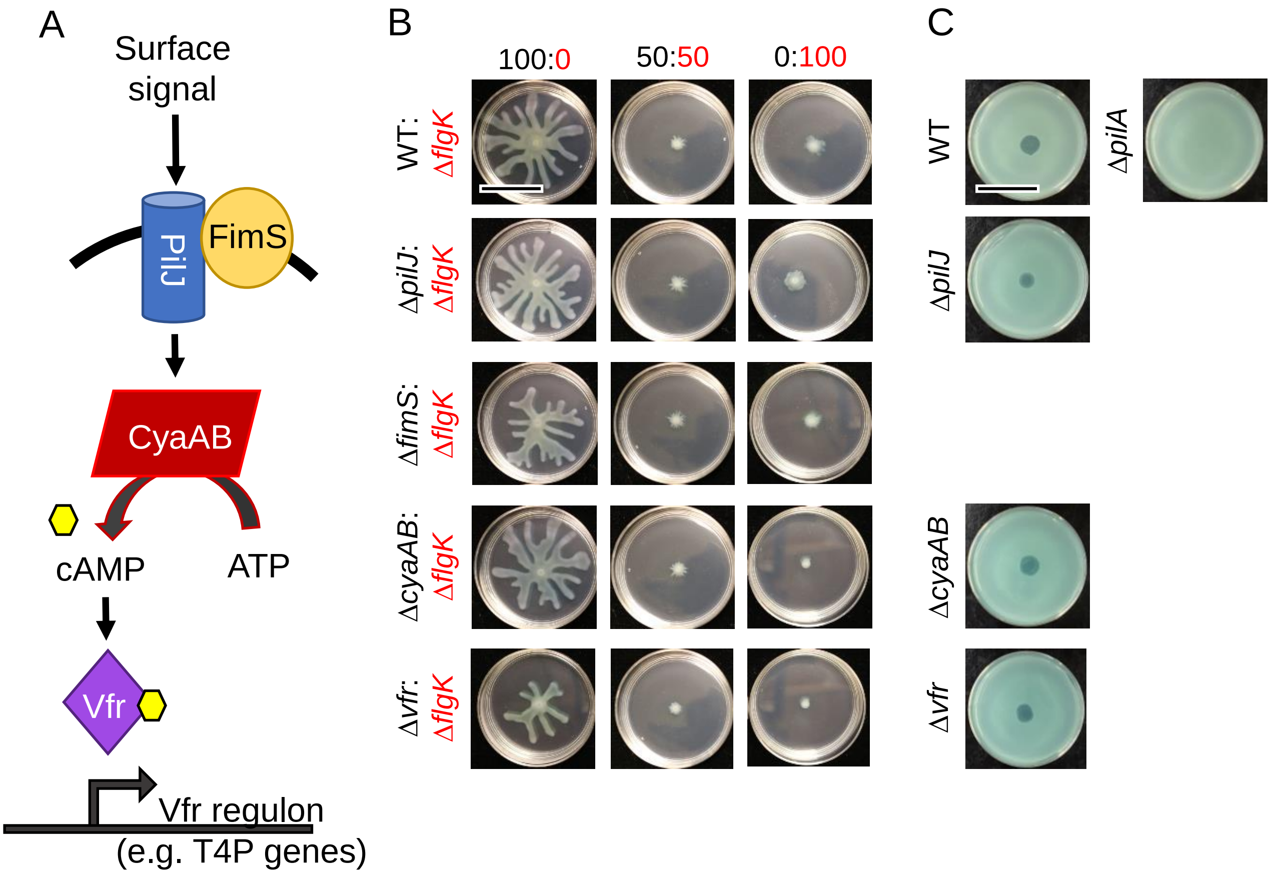


**Fig. S7. Surface-sensing and cAMP signaling is not involved in mixed motility repression. A.** Schematic of genes involved in surface sensing in *P. aeruginosa*. This pathway is dependent on the perception of a surface followed by cAMP induction and transcriptional activation of the Vfr regulon. Schematic was adapted from Luo *et al.,* 2015. **B.** Representative images from swarm assays of wild-type (WT) *P. aeruginosa* strain PA14, Δ*flgK*, Δ*pilJ,* Δ*fimS,* Δ*cyaAB,* and Δ*vfr* mixed at the indicated combinations and ratios on M8 agar. **C.** Representative images from plaque assays of WT, Δ*pilJ,* Δ*cyaAB,* Δ*vfr*, and Δ*pilA* (negative control) showing sensitivity (T4P+) and insensitivity (T4P-) to phage DMS3 lysis. Data are representative of three individual experiments. WT and derivatives are labeled in black while Δ*flgK* is labeled in red. Scale bar: 30 mm

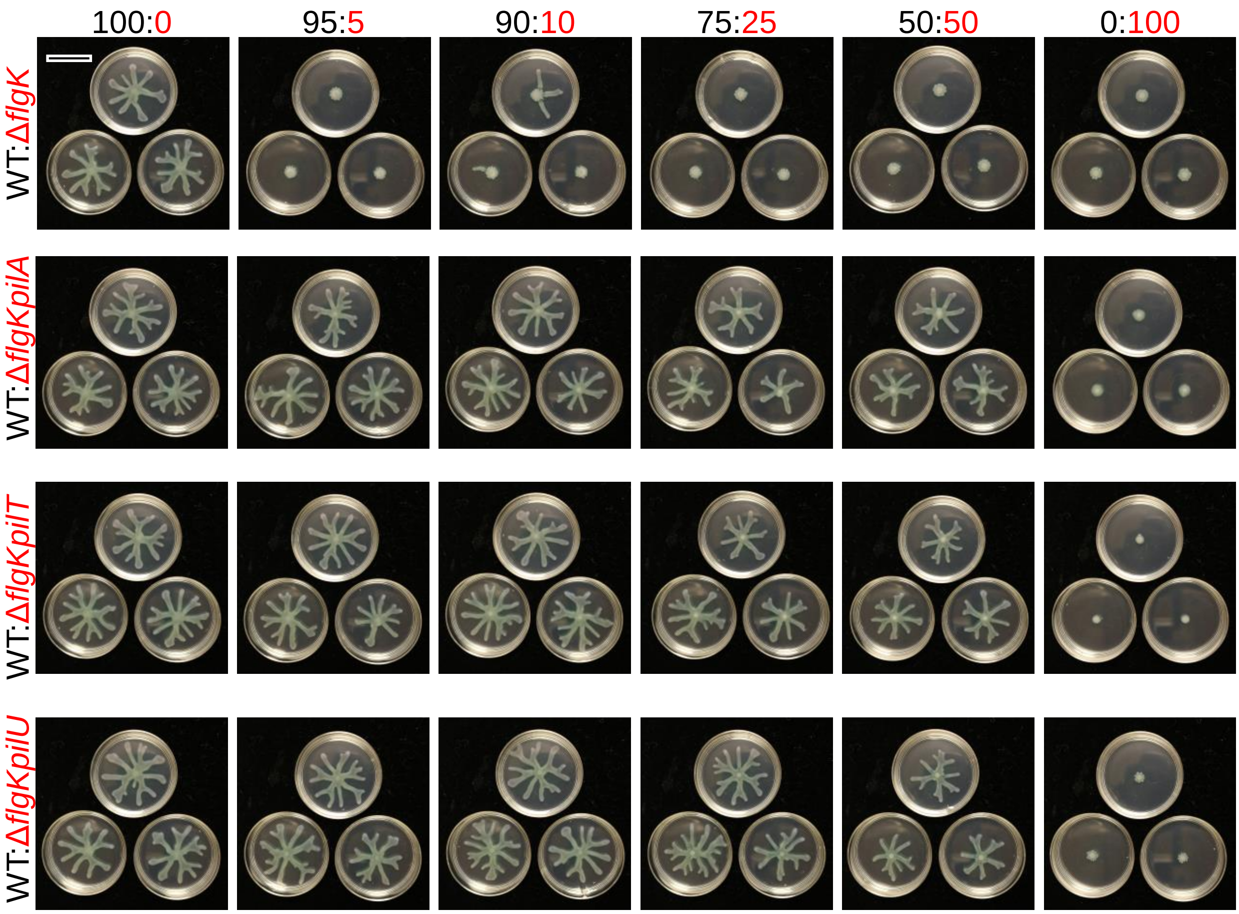


**Fig. S8. Non-flagellated *P. aeruginosa* requires functional T4P to repress swarming by flagellated strains on soft agar.** Representative images from swarm assays of wild-type (WT) *P. aeruginosa* strain PA14 (flagellated and T4P+) mixed with Δ*flgK* (non-flagellated and T4P+), Δ*flgK*Δ*pilA* (non-flagellated and T4P-), Δ*flgK*Δ*pilT* (non-flagellated with no T4P retraction), or Δ*flgK*Δ*pilU* (non-flagellated with reduced T4P retraction) at ratios indicated. Images show three technical replicates from a single experiment. Images are representative of three individual experiments. WT and derivatives are labeled in black while Δ*flgK* is labeled in red. Scale bar: 30 mm

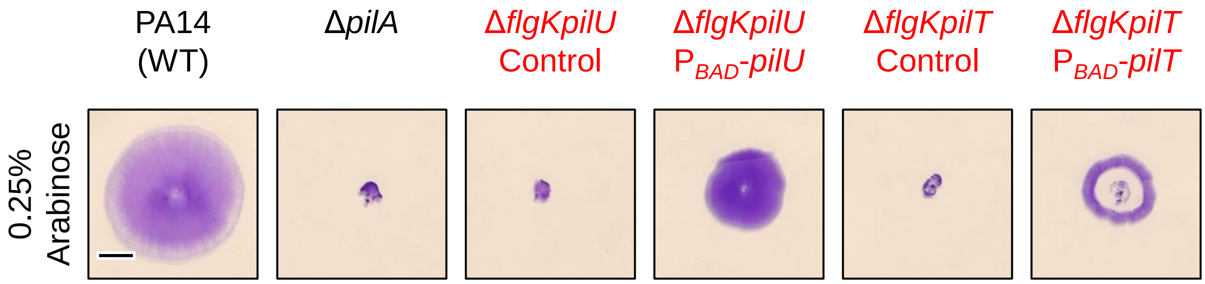


**Fig. S9. Complementation of Δ*flgKpilU* and Δ*flgKpilT***. Representative images from twitch assays of wild-type (WT) *P. aeruginosa* strain PA14, Δ*pilA,* Δ*flgKpilU,* Δ*flgKpilU* $P_{BAD}$-*pilU* (complemented), Δ*flgKpilT*, and Δ*flgKpilT* $P_{BAD}$-*pilT* (complemented) on T-broth agar containing 0.25% arabinose after 46 h incubation at 37°C. Data are representative of three individual experiments. Scale bar: 5 mm

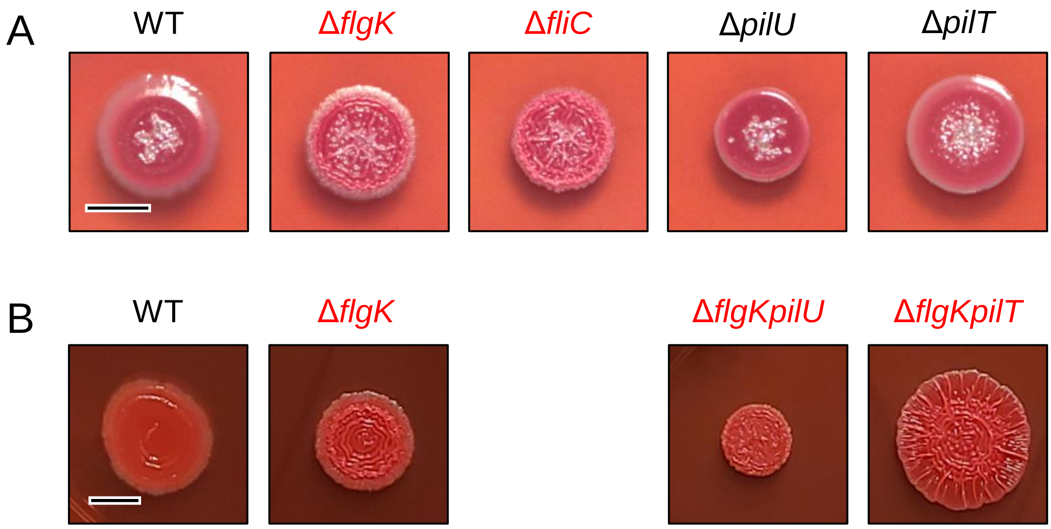


**Fig. S10. Non-flagellated mutants have increased Congo Red binding indicative of increased Pel production.** Representative images of **A.** wild-type (WT) *P. aeruginosa* strain PA14 (flagellated and T4P+), Δ*flgK* (non-flagellated and T4P+), Δ*fliC* (non-flagellated and T4P+), Δ*pilU* (flagellated with reduced T4P retraction), Δ*pilT* (flagellated with no T4P retraction) and **B.** WT, Δ*flgK,* Δ*flgK*Δ*pilU* (non-flagellated with reduced T4P retraction), and Δ*flgK*Δ*pilT* (non-flagellated with no T4P retraction) from two separate experiments grown on Congo Red plates to assess Pel production. WT and derivatives are labeled in black while Δ*flgK* and derivatives are labeled in red. Scale bar: 5 mm
